# *MyD88* deficiency modestly attenuates disease in a Leigh syndrome mouse model while enrofloxacin accelerates disease

**DOI:** 10.64898/2026.05.13.724988

**Authors:** Allison R Hanaford, Elizaveta A Olkhova, Ryan Liao, Ashley Ching, Alvin Huang, Erin Shien Hsieh, Kino Watanabe, Yihan Chen, Megan Wichman, Noelle Hwang, Katerina James, Michael Mulholland, Vivian Truong, Holly Coulson, Kerry Gibbins, Owen Cairns, Anastasia Dimitriou, Bernhard Kayser, Brittany M Johnson, Surojit Sarkar, Vandana Kalia, Simon C Johnson

**Affiliations:** Ben Towne Center for Childhood Cancer and Blood Disorders Research, Seattle Children’s Research Institute, Seattle WA USA; Translational Bioscience, Northumbria University, Newcastle, UK; Norcliffe Foundation Centre for Integrative Brain Research, Seattle Children’s Research Institute, Seattle, WA, USA; Center for Integrated Brain Research, Seattle Children’s Research Institute, Seattle WA USA; Department of Pediatrics, University of Washington, School of Medicine

**Keywords:** Leigh syndrome, mitochondrial disease, innate immune system, electron transport chain complex I

## Abstract

Primary genetic mitochondrial diseases (GMDs) are a clinically and genetically diverse group of diseases estimated to impact over 1 in 4,000 individuals. Leigh syndrome (LS) is the most common pediatric presentation of GMD. LS typically presents within the first years of life and is a severe progressive multi-system disorder. Symmetric progressive inflammatory brain lesions are a defining feature of the disease. Patients can also present with seizures, metabolic dysfunction, muscle weakness, and other symptoms. No effective clinical treatments currently exist. Recent data from the *Ndufs4*(-/-) mouse model shows that peripheral macrophages contribute to brain lesions in LS, that disease is causally driven by innate immune populations, and that depletion of innate immune cells prevents LS disease. However, the precise mechanisms underlying immune activation remain unknown. Certain mitochondrial macromolecules retain bacterial signatures and can act as potent agonists for innate immune pathways. For example, cytoplasmic mitochondrial RNA and mitochondrial DNA are detected by Toll-like receptors (TLRs) 7 and 9, respectively, at the endosome. Accordingly, these are considered strong candidates for mediating innate immune activation in LS. Here, we generated TLR signaling deficient *Ndufs4*(-/-)/*MyD88*(-/-) animals to assess whether TLR signaling plays a role in disease onset or progression in LS. Loss of *MyD88* in *Ndufs4*(-/-) animals statistically significantly increased survival and delayed the onset of some symptoms, but the benefits were modest compared to CSF1R inhibition from prior work. We conclude that *Myd88*-mediated immune signaling is not a primary driver of LS. Notably, prophylactic enrofloxacin treatment, which was necessary for production of innate immune deficient *MyD88*(-/-) animals, modestly decreased survival and accelerated disease. The impact of enrofloxacin and similar drugs in the context of mitochondrial disease warrants further investigation.

## Introduction

Genetic mitochondrial diseases (GMDs) are a genetically and clinically diverse collection of diseases estimated to impact over 1 in 4,000 individuals. Leigh syndrome (LS) is the most common pediatric presentation of GMD. Symmetric progressive necrotizing lesions in specific brain regions are a defining clinical feature of LS. Patients can also experience seizures, metabolic dysfunction, muscle weakness, ataxia, other symptoms. Over 120 individual unique genes have been causally associated with LS (1). The precise mechanisms of disease pathobiology in LS remain unclear, and no effective clinical interventions are available. The extreme genetic diversity limits the therapeutic potential of gene-specific interventions such as gene therapy, emphasizing the need for research into mechanisms of disease in order to find common targets for treatment.

While effective clinical therapeutics are lacking, recent studies have identified promising treatment strategies which robustly prevent or reverse disease in animal models. In the *Ndufs4*(-/-) mouse model of LS, considered the premier model of GMD due to the close resemblance to severe forms of human LS, we have shown that immune depletion prevents LS symptoms (2). The adaptive immune system appears to play no role, with CSF1R (colony stimulating factor-1) dependent innate immune cells apparently driving the full disease (2, 3). Separately, we’ve shown that peripheral immune cells invade the brain to drive CNS lesions in coordination with brain resident microglia, and that peripheral immune cells are sufficient to cause the full disease in mice lacking microglia (4).

While we have now established that immune cell actions play a causal role in LS, the precise mechanisms underlying immune activation remain unknown. Certain mitochondrial macromolecules retain bacterial signatures and can act as potent agonists for innate immune receptors (5). It is already established that mitochondrial components can act as damage associated molecular patterns (DAMPs) or be recognized as pathogen associated molecular patterns (PAMPs) due to their similarity to bacterial or viral components (5-7) . These DAMPs and PAMPs are recognized by pattern recognition receptors (PRRs) and induce innate immune responses. The possibility that these factors somehow leak or are actively exported into the cytoplasm or extracellular spaces and activate in LS is an attractive hypothesis for activation of the immune system in LS.

One major class of PRRs involved in recognizing mitochondrial macromolecules as DAMPs or PAMPs are the Toll-Like Receptors (TLRs). The TLRs are an ancient and highly conserved family of receptors widely present in vertebrates, invertebrates, and even plants (8). Humans have at least 10 TLRs, mice at least 12 (9). These are typically highly expressed in dendritic cells and macrophages/monocytes (10). Some mammalian TLRs localize to the plasma membrane while others localize to endosomes and detect internalized materials (10). Key mitochondrial components known to activate TLRs include mitochondrial DNA (TLR9) (6, 7), mitochondrial RNA (TLR7) (11), and heat shock proteins (TLR4) (12) (see ***Discussion*** for more details). Accordingly, TLRs are considered strong candidates for mediating innate immune activation in LS.

While the large number of TLRs presents challenges for systematic study, the majority of TLRs (excluding TLR3) act through the adapter protein MyD88 (Myeloid differentiation primary response 88), which also acts downstream of IL-1Rs. Although some non-MyD88 pathways also exist (see ***Discussion***), genetic disruption of *MyD88* allows for a broad assessment of the role of TLR signaling in a given phenotype.

Here, we generated TLR signaling deficient *Ndufs4*(-/-)/*MyD88*(-/-) animals to assess whether TLR signaling plays a role in disease onset or progression in LS.

## Methods

### Animals

*MyD88*(-/-) mice were obtained from the Jackson Lab (strain # 009088) and were originally developed by Hou et al (13). *Ndufs4*(+/-) mice were originally developed and obtained from the Palmiter laboratory at the University of Washington, Seattle WA (14) and are available at the Jackson laboratory (Strain #027058). Genotyping of *Ndufs4* and *MyD88* was performed according to the Jackson Laboratory methods (for strains #009088 and #027058). Only mice with PCR confirmed genotypes were used in the study. *Ndufs4*(-/-) mice cannot breed due to disease pathology and short lifespan, so all *Ndufs4*(-/-) mice were generated using *Ndufs4*(+/-) breeders.

*Myd88*(-/-) mice were crossed with *Ndufs4*(+/-) animals to generated *Ndufs4*(+/-)/*MyD88*(+/-) breeders. As discussed in detail in **Results**, we initially failed to generate *MyD88*(-/-) animals from *Ndufs4*(+/-)/*MyD88*(+/-) crosses. Veterinary laboratory testing confirmed the presence of common mouse pathogens not excluded from specific pathogen free (SPF) housing at Seattle Children’s Research Institute (SCRI) where the mice were housed. *Ndufs4*(+/-)/*MyD88*(+/-) breeders and offspring were subsequently provided prophylactic antibiotic enrofloxacin delivered in the drinking water which led to *MyD88*(-/-) pups being born at the expected ratio. Breeders and experimental mice were maintained on ∼85 mg/kg/day of enrofloxacin (based on water intake) for the duration of the study. Medicated water was changed weekly.

Mice were weaned at 20-22 days of age. To promote normal body temperature and activity, *Ndufs4*(-/-) mice were always housed with control littermates for warmth, as in our prior work (2, 3, 14-19). Mice were weighed and health assessed a minimum of 3 times a week from weaning to P35 (the approximate age of disease onset) and 5 times a week from P35 until the end of the study. Following onset of symptoms in *Ndufs4*(-/-) mice, wet food was provided at the bottom of the cage. Animals were euthanized if the any of the following criteria were met: loss of 20% of body weight, from individual mouse maximum, measured on two consecutive days; immobility; or any signs of severe acute distress. *Ndufs4*(+/-) have no detected phenotype, so controls consist of both *Ndufs4*(+/-) and *Ndufs4*(+/+) mice.

Clasping and ataxia were assessed by visual scoring as in our previous studies (4, 15, 20-22). As *Ndufs4*(-/-) mice can show transient/intermittent improvement of symptoms during disease progression, for all disease symptoms we report if an animal has ever displayed a particular symptom for two or more consecutive days. Onset of weight is reported as age of maximum body weight based on final weight curves at the end of the study.

Experimental mice were fed Picolab diet 5058 and were housed on a 12-hour light cycle. Breeders wer fed PicoLab diet 5053. Animal experiments followed SCRI guidelines and were approved by the SCRI IACUC. In these and our prior studies runts (defined as ≤5 g body weight at weaning age), those that experienced weaning stress, or those or born with health issues unrelated to the *Ndufs4*(-/-) phenotype (such as hydrocephalus) were euthanized and excluded at or prior to weaning. These exclusions are uncommon (estimated to occur approximately once per 5-10 litters), were not increased in this study versus prior studies, and appear to be unrelated to the genes of interest for these studies (see ***Table 1*** and ***Table 2***).

**Table 1.** Offspring ratios from *Ndufs4*(+/-)/*MyD88*(+/-) breeding pairs. Without enrofloxacin, *MyD88*(-/-) mice were produced at a significantly lower than expected rate (X^2^ p-value ****9.52×10^-7^), with only one born out of a total of 116 pups.

Without Enrofloxacin
| Possible Genotype | Expected Frequency | Expected Numbers | Observed Number |
| --- | --- | --- | --- |
| *Ndufs4*(+/-)/*MyD88*(+/-) X *Ndufs4*(+/-)/*MyD88*(+/-) |  |  |  |
| Total pups: 116 |  |  |  |
| *Ndufs4*(+/-)/*MyD88*(+/-) | 0.25 | 29 | 34 |
| *Ndufs4*(+ +)/*MyD88*(+/-) | 0.125 | 14.5 | 24 |
| *Ndufs4*(+/-)/*MyD88*(+ +) | 0.125 | 14.5 | 23 |
| ***Ndufs4*(+/-)/*MyD88*(-/-)** | **0.125** | **14.5** | **1** |
| *Ndufs4*(-/-)/*MyD88*(+/-) | 0.125 | 14.5 | 13 |
| ***Ndufs4*(+ +)/*MyD88*(-/-)** | **0.063** | **7.308** | **0** |
| *Ndufs4*(+ +)/*MyD88*(+ +) | 0.063 | 7.308 | 9 |
| *Ndufs4*(-/-)/*MyD88*(+ +) | 0.063 | 7.308 | 12 |
| ***Ndufs4*(-/-)/*MyD88*(-/-)** | **0.063** | **7.308** | **0** |

$\chi^2$ p-value\*\*\*\*: $9.52 \times 10^{-7}$

**Table 2.** Offspring ratios from *Ndufs4*(+/-)/*MyD88*(+/-) breeding pairs treated with prophylactic Enrofloxacin. After beginning prophylactic treatment of colony with enrofloxacin the rate of *MyD88*(-/-) pup production rose to expected rates. Following the successful births of those animals represented in this table we bred animals using *Ndufs4*(+/-)*/MyD88*(-/-) breeders.

With Enrofloxacin
| Possible genotype | Expected Frequency | Expected Numbers | Observed Number |
| --- | --- | --- | --- |
| *Ndufs4*(+/-)/*MyD88*(+/-) X *Ndufs4*(+/-)/*MyD88*(+/-) |  |  |  |
| Total pups: 34 |  |  |  |
| *Ndufs4*(+/-)/*MyD88*(+/-) | 0.25 | 8.5 | 12 |
| *Ndufs4*(+ +)/*MyD88*(+/-) | 0.125 | 4.25 | 1 |
| *Ndufs4*(+/-)/*MyD88*(+ +) | 0.125 | 4.25 | 5 |
| ***Ndufs4*(+/-)/*MyD88*(-/-)** | **0.125** | **4.25** | **1** |
| *Ndufs4*(-/-)/*MyD88*(+/-) | 0.125 | 4.25 | 3 |
| ***Ndufs4*(+ +)/*MyD88*(-/-)** | **0.063** | **2.142** | **3** |
| *Ndufs4*(+ +)/*MyD88*(+ +) | 0.063 | 2.142 | 3 |
| *Ndufs4*(-/-)/*MyD88*(+ +) | 0.063 | 2.142 | 4 |
| ***Ndufs4*(-/-)/*MyD88*(-/-)** | **0.063** | **2.142** | **2** |

$\chi^2$ p-value: 0.32 (n.s.)

### Nanostring and qPCR

Flash-frozen brain tissue samples were homogenized using Dounce homogenizers. Each frozen brainstem tissue sample was dropped into 1 mL of frozen QIAzol solution in the dounce homogenizer and crushed into the QIAzol using the homogenizer. The solution was transferred to an Eppendorf tube and placed on dry ice, and samples were stored at -80° C until RNA extraction using Qiagen RNeasy Lipid Tissue Mini kit following manufacturer’s guidelines. RNA concentration and quality were assessed using NanoDrop 2000.

RNA used for NanoString analysis was further assessed using an agarose-bleach gel method, as detailed in (23). Only samples with acceptable 260/230 and 260/280 ratios and no evidence of degradation by gel analysis were used for the NanoString assay. These were serially diluted, with concentration checked at the 10X concentration, to reach a final concentration of 20 ng/ µL, of which 5 µL each was submitted for analysis. Brainstem gene expression analysis was assessed using NanoString mouse cancer immune panel (NS_Mm_CancerImm_C3400), which was analyzed by the Genomics Resource Center at the Fred Hutchinson Cancer Research Center. Transcript counts were normalized to *Eefg1*, the housekeeping gene showing the lowest variance between treatment groups and are presented as normalized counts (versus control animals).

For qPCR, 1500 ng of RNA was used as a template for reverse transcription utilizing SuperScript VILO cDNA Synthesis Kit (Invitrogen; catalogue #11754-050) following manufacturer’s instructions. qRT-PCR was performed using Universal TaqMan MasterMix and TaqMan FAM probes against the genes of interest, as per manufacturer recommendations. Probes used were: *Tlr7* - Mm00446590_m1 (FAM-MGB); *Tlr9* - Mm00446193_m1 (FAM-MGB); *MyD88* - Mm00440338_m1 (FAM-MGB); *Actin* - Mm02619580_g1 (VIC-MGB). All targets of interest were labelled with FAM and run together with the VIC labelled probe against actin (*ActB*) to which all the data were normalized. A relative standard curve was included in every run and used to calculate relative expression.

### Tissue Collection and immunofluorescent staining

Tissue was collected when euthanasia criteria were met. Mice were fully anaesthetized and perfused with approximately 20 mL of chilled PBS followed by approximately 20 mL of chilled 4% PFA. Immediately following perfusion, the brain was removed and placed in ∼20mL 4% PFA for a minimum of 24 hours at 4º C. The brain was then moved to cryoprotection solution (30% sucrose, 1% DMSO, 100 µM glycine, 1XPBS) for a minimum of 48h at 4º C. Cryopreserved tissue was frozen in Tissue-Tek O.C.T. compound in cryoblocks on dry ice and stored at -80º C until cryo-sectioning. Cryoblocks were sectioned at 50 μm thickness using a Leica CM30505 cryostat set at -20º C. Slices were immediately placed in 1XPBS with 1 µg/mL DAPI at 4º C for a maximum of 24 hours before mounting on slides. Prior to mounting, slices were examined for the presence of lesions using a fluorescent microscope. Slices with lesions were mounted on SuperFrost Plus slides. Sections from the center of the lesion were chosen for staining and are shown here. Mounted sections were baked at 37º C in a white-light LED illuminated incubator for 24 hours and then storing at -80º C until staining.

For immunofluorescent staining, slides were incubated for 24 hours at 50º C in antigen retrieval buffer (10 mM citrate, pH 6.0). Following antigen retrieval, slides were washed for 5 minutes in 1X PBS on ice before being incubated for 1 hour on ice in 1 mg/mL sodium borohydride in 1XPBS. Slides were then washed in 1X PBS with 10 mM glycine for 5 minutes before being placed in a 0.5 mg/mL Sudan Black in 70% ethanol and incubated overnight with gentle stirring. Slides were then washed 3X for 5 minutes each with 1X PBS. Excess moisture was wiped away and a hybriwell sealing sticker (Grace Bio-Labs, GBL612202) was applied to each slide over the tissue slices. Slides were incubated for 30 minutes at room temperature in blocking/permeabilization solution (1 mM digitonin, 10% rabbit serum, 0.5% tween in 1X PBS). Following blocking, the hybriwell sealing stickers were removed and the blocking solution was shaken off the slides. Excess moisture was wiped away and new hybriwells were placed. The slides were incubated for 72 h hours at 4º C in conjugated primary antibodies and 1 µg/mL DAPI diluted in the blocking solution. The following fluorescently conjugated primary antibodies were used: Anti-Iba1 (1:150, Wako 013-26471), anti-CD45 (1:150, clone D3F8Q, Cell Signaling Technologies #19581). After incubation slides were washed for 5 minutes 4X in 1X PBS. ProLong Gold Antifade was used as mounting media and coverslips sealed with clear nail polish. Slides were stored in an opaque slide box at 4º C until imaging.

Slides were imaged using a Zeiss LSM900 microscope using an EC Plan-Neoflar 10x/0.3 M27 objective. Channels were set to an optical thickness of 12.5 microns. DAPI was excited with a 405nm laser, Alexa488 (Cd45) with a 488nm laser, and Alex635 (Iba1) with a 633nm laser. Images were acquired using approximately the same laser intensity. Images were post-processed using the ZEISS ZEN Microscopy Software (RRID:SCR_013672) LSM Plus module which uses a Wiener filter to enhance signal-noise ratio (SNR), improve resolution and provide spatial information. Brightness/Contrast were adjusted in Fiji (24) for visualization to make cell morphology visible.

### Statistical Analysis

All statistical analyses were performed using GraphPad Prism 10.6.1 as detailed in figure legends.

### Scientific rigor

Sex-Both male and female mice are used in these experiments. No differences between male and female *Ndufs4*(-/-) or *Myd88*(-/-) mice have been reported or observed by us.

## Results

### TLRs expression is upregulated in Ndufs4(-/-) mouse brainstem lysates

Prior studies have demonstrated peripheral immune cells invade the CNS and drive lesions and elimination of immune cells prevents disease in the *Ndufs4*(-/-) mouse model of Leigh syndrome (2, 25). However, disrupting the adaptive immune system alone by depleting B-, T-, and NK-cells has no effect on disease (3). In an effort to identify innate immune pathways that may be contributing to disease pathogenesis we performed a targeted analysis of immune-related gene expression in *Ndufs4*(-/-) and control mouse brainstem lysates using the NanoString cancer immune panel. The majority of TLRs show increased expression in the *Ndufs4*(-/-) brainstem, with significant increases in TLRs 2, 3, 5, 7, and 9 (***Fig. 1A***). There was no significant change in *MyD88* expression. Changes in expression of *Tlr7, Tlr9*, and *Myd88* were confirmed by qPCR analysis of brainstem lysates from additional animals (***Fig. 1B***).

**Figure 1.**
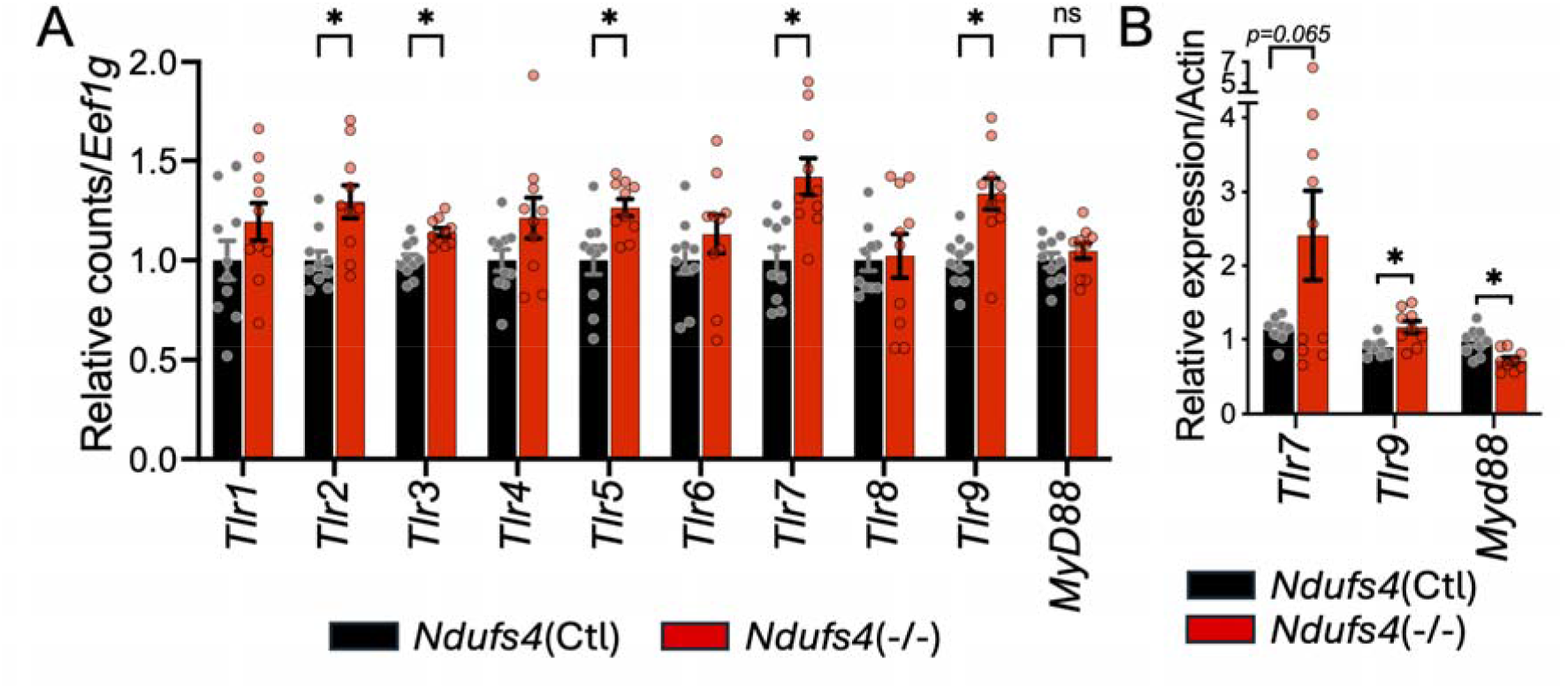
Toll-like receptor expression is increased in the brainstem of diseased *Ndufs4*(-/-) mice. A) NanoString-based analysis of Toll-like receptor (*Tlr*) and *MyD88* expression in the brainstem of 45-day old control and *Ndufs4*(-/-) mice (see ***Methods***). Black – *Ndufs4*(Ctl), Red – *Ndufs4*(-/-) (see ***Methods***). p-values reflect multiple testing corrected (Holm-Šídák method) pairwise t-tests. *p<0.05, ns and those not shown – not significant. *B*) Taqman probe-based qPCR analysis of *Tlr7, Tlr9*, and *MyD88* expression in 50-day old control and *Ndufs4*(-/-) mouse brainstem samples. Normalized to actin, included in each reaction. Black – *Ndufs4*(Ctl), Red – *Ndufs4*(-/-) (see ***Methods***). Relative expression calculated using relative standard curve approach. *p<0.05, ns – not significant, by Holm-Šídák method multiple-testing corrected pairwise t-tests with Welch’s correction (no assumption regarding standard deviation).

### Generation of Ndufs4(-/-)/MyD88(-/-) mice and prophylactic enrofloxacin

To determine the role of MyD88 dependent TLR signaling in the pathogenesis of LS, we crossed *Ndufs4*(-/-) and *MyD88*(-/-) animals to generate *Ndufs4*(+/-)/*MyD88*(+/-) animals to use to as breeders to produce *Ndufs4*(-/-)/*MyD88*(-/-) double knockout mice. During our first months of breeding *Ndufs4*(+/-)/*MyD88*(+/-) x *Ndufs4*(+/-)/*MyD88*(+/-) mice we did not detect any *MyD88* knockout animals. Specifically, of 105 genotyped pups from *Ndufs4*(+/-)/*MyD88*(+/-) crosses, only one *MyD88*(-/-) pup was born, significantly lower than expected (see ***Table 1***). *MyD88* deficiency is known to result in compromised innate immune responses, a fact central to our study design (13, 26). Given that our mice were breeding in a specific pathogen free (SPF) environment, we considered the possibility that a non-excluded pathogen present in our colony was leading to loss of *MyD88*(-/-) pups in utero or soon after birth (prior to weaning and genotyping).

Veterinary PCR testing of breeder feces and saliva (IDEXX) identified common mouse pathogens in our colony, including two species of *Helicobacter* detected in faces (see ***Supplemental Data***). Immune-compromised mice have been shown to be susceptible to *Helicobacter*, and there is evidence that *Helicobacter* impacts pregnancy outcomes in mice including increasing fetal absorption (27). Accordingly, we provided prophylactic enrofloxacin, a commonly used veterinary antibiotic, via water bottles at ∼85 mg/kg/day of enrofloxacin (based on water intake). In breeders treated and maintained on enrofloxacin *MyD88*(-/-) mice were produced at expected ratios (see ***Table 2***). Following the successful production of *MyD88*(-/-) animals with administration of enrofloxacin, we also bred experimental mice using *Ndufs4*(+/-)/*MyD88*(-/-) breeders.

*MyD88* deficiency is known to be autosomal recessive in humans and mice, with one copy of MyD88 sufficient for normal signaling ((28, 29), see ***Methods***, Jackson Laboratory strain #009088 information). In agreement, we observed no differences between *MyD88*(+/-) and *MyD88*(+/+) animals in either treatment group, so *MyD88*(+/-) and *MyD88*(+/+) were pooled as *MyD88*(Ctl) for analysis (raw data provided in supplemental files).

While effective in allowing for the generation of *MyD88*(-/-) animals, antibiotics in general, and enrofloxacin specifically, have been reported to impact some mitochondrial functions (see ***Discussion***, (30, 31)). Accordingly, a non-enrofloxacin treated cohort is included for comparison, *Ndufs4*(-/-)/*MyD88*(-/-) animals could only be produced with enrofloxacin, so are only compared against a similarly enrofloxacin treated *MyD88* control group.

### MyD88 deficiency does not significantly impact maximum weight or onset of cachexia

Weight in *Ndufs4*(-/-) animals reaches a maximum around P37 followed by a disease-associated loss of weight until death (2, 3, 18). Overall weight trajectory and maximum weight were not significantly altered by either *MyD88* loss or enrofloxacin treatment (***Fig. 2A-B***). Neither *MyD88* deficiency or enrofloxacin had a significant impact on the age of onset of cachexia (weight loss) (***Fig. 2C***).

**Figure 2.**
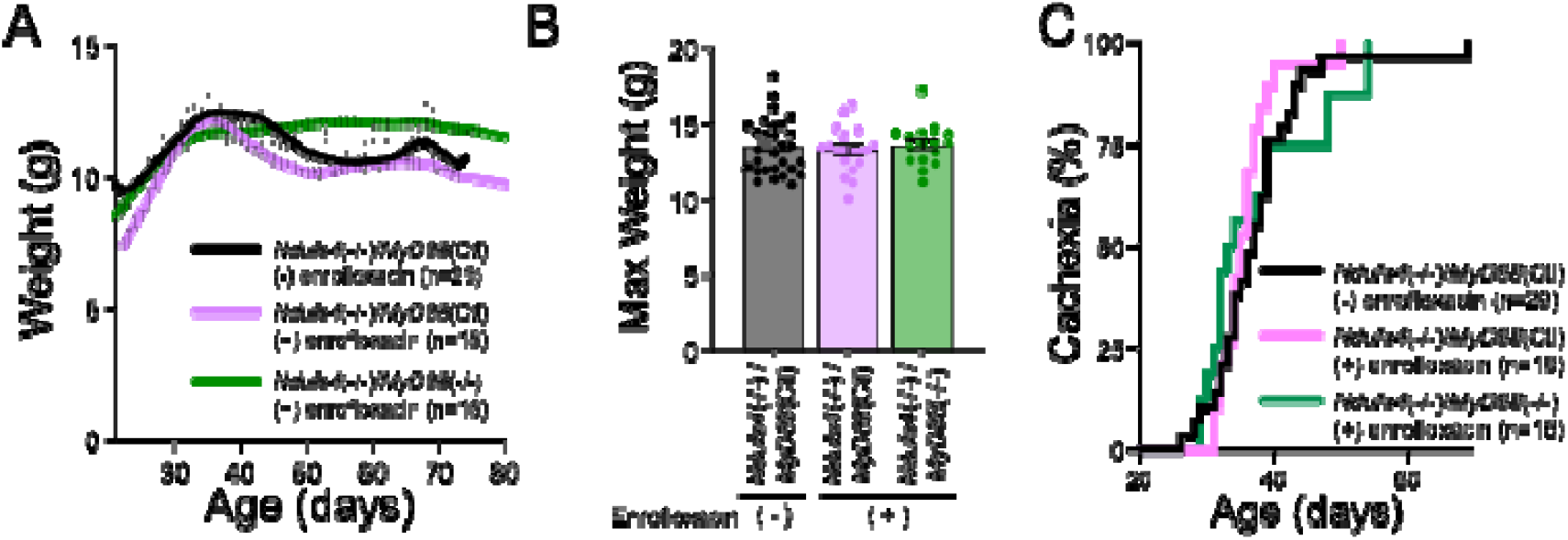
*MyD88* deficiency and enrofloxacin have no significant effect on cachexia onset or maximum weight in *Ndufs4*(-/-) mice. A) Impact of enrofloxacin and *MyD88* disruption on overall weight trajectory in *Nduf4*(-/-) mice (see ***Results, Table 1, 2***). Replicates (n’s) as indicated. B) Maximum weights of individual animals in A). Replicates (n’s) as indicated in A). ANOVA and pairwise comparisons were not statistically significant. C) Impact of enrofloxacin and *MyD88* disruption on the onset of cachexia (weight loss) in *Nduf4*(-/-) mice. No curves significantly different by pairwise log-rank comparison. Replicates (n’s) as indicated.

### Enrofloxacin and MyD88 deficiency differentially impact neurologic disease onset

In addition to progressive weight loss, *Ndufs4*(-/-) mice present with neurologic symptoms including ataxia and forelimb clasping, a behaviorally assessed sign of degenerative neurologic disease in mice (32). As with weight loss, onset of neurological symptoms occurs around P37 in *Ndufs4*(-/-) animals. Enrofloxacin was associated with a significantly earlier presentation of ataxia when comparing *Ndufs4*(-/-)/*MyD88*(Ctl) mice with and without the drug (***Fig. 3A***, *see* ***Discussion***). Loss of *MyD88* did not significantly delay the onset of ataxia when comparing enrofloxacin treated *Ndufs4*(-/-)/*MyD88*(Ctl) and *Ndufs4*(-/-)/*MyD88*(-/-) animals. Onset of forelimb clasping was significantly (but modestly) delayed by loss of *MyD88* (***Fig. 3B***). Enrofloxacin treatment was associated with an earlier onset of forelimb clasping, but this was not statistically significant.

**Figure 3.**
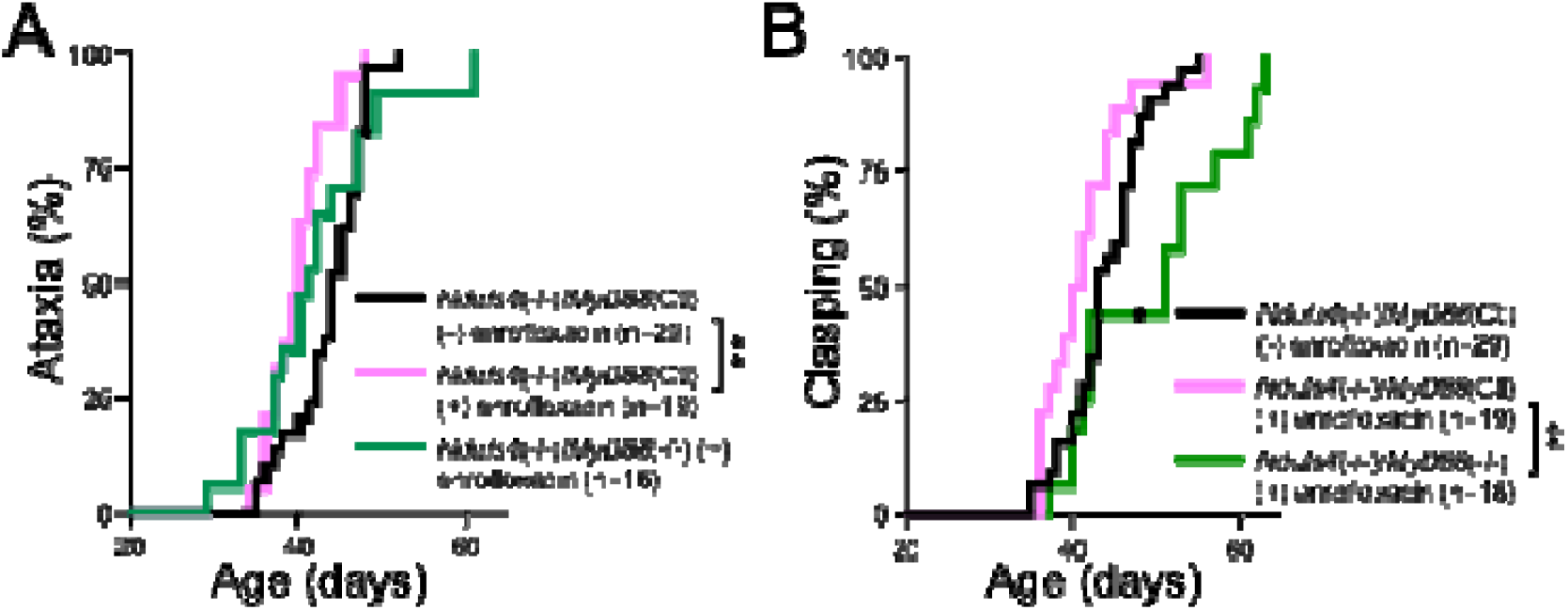
*MyD88* deficiency and enrofloxacin modestly impact neurologic disease onset in *Ndufs4*(-/-) mice. A) Impact of enrofloxacin and *MyD88* disruption on ataxia onset in *Nduf4*(-/-) mice. n’s as indicated. The dot (MyD88(-/-) cohort) represents an animal which died prior to presenting with ataxia. P-value shown indicates pairwise log-rank test comparisons between curves indicated - **p<0.005. B) Impact of enrofloxacin and *MyD88* disruption on the onset of clasping in *Nduf4*(-/-) mice. n’s as indicated. The dot (MyD88(-/-) cohort) represents an animal which died prior to presenting with clasping. P-value shown indicates pairwise log-rank test comparisons between curves indicated - **p<0.005.

### MyD88 deficiency did not alter CNS lesion formation

Loss of *MyD88* did not prevent overt neuroinflammation, including brainstem lesions and inflammation in the cerebellar peduncle, in *Ndufs4*(-/-) animals as assessed by immunofluorescent staining for microglia/macrophages with Iba1 (ionized calcium-binding adapter molecule 1) and peripheral immune cells (CD45) (***Fig. 4***, see (25), ***Discussion***). Lesions are similarly present in all enrofloxacin groups (not shown).

**Figure 4.**
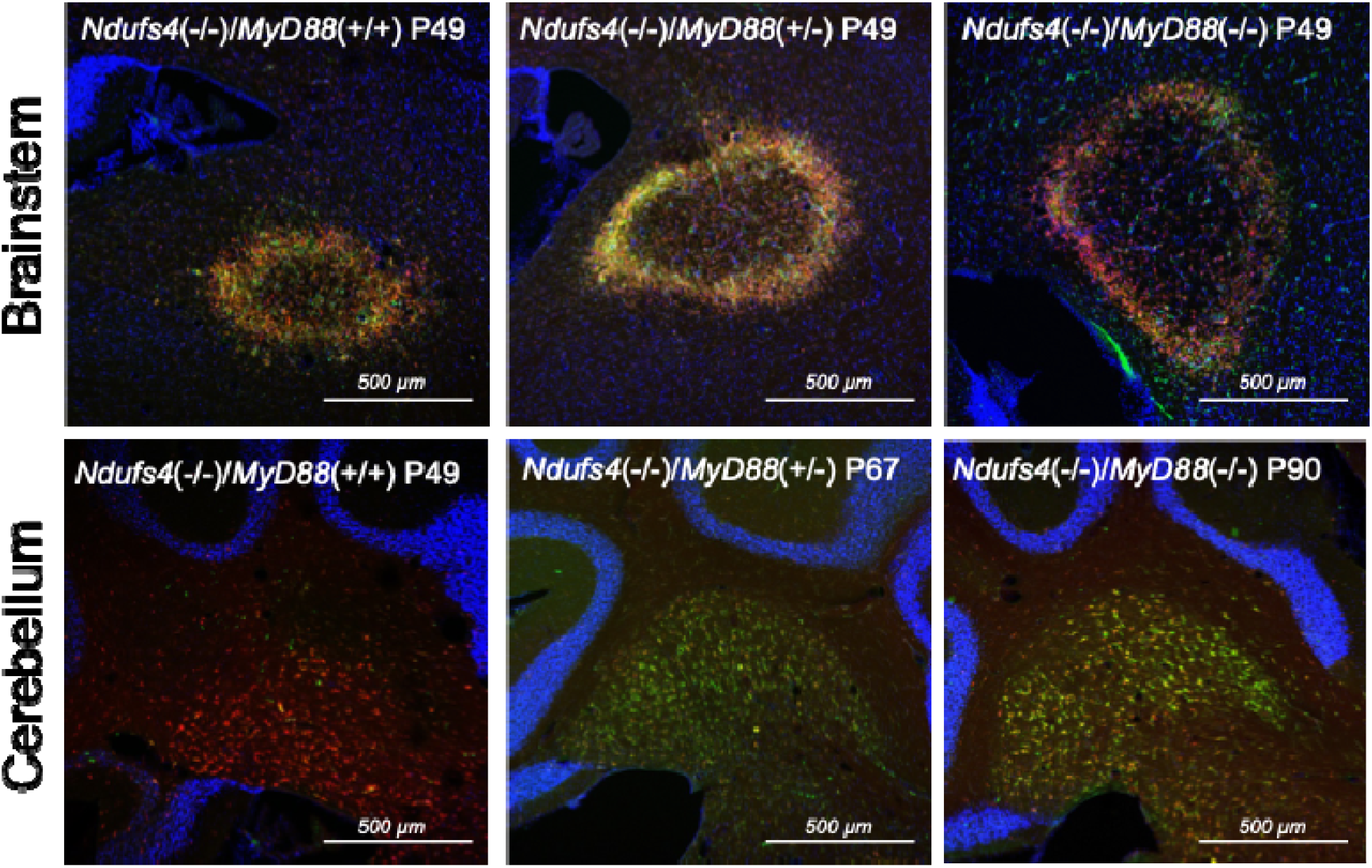
*MyD88* deficiency does not prevent CNS lesions in *Ndufs4*(-/-) mice. Immunofluorescent staining of brainstem and cerebellar sites of neuroinflammation in the *Ndufs4*(-/-) model, with brainstem slices focused on neuroinflammatory lesion sites. Red – Iba1 (ionized calcium-binding adapter molecule 1), green – CD45, blue – DAPI (4’,6-diamidino-2-phenylindole)(see ***Methods***). Size of image fields - 1413.19 µm x 1413.19 µm, scalebar ∼500 µm.

### Enrofloxacin and MyD88 deficiency have modest opposing effects on median survival

Disease progression is rapid in untreated *Ndufs4*(-/-) mice with a median survival of ∼50-60 days. The most frequent cause of death is euthanasia due to reaching a pre-determined weight-loss threshold (20% loss from maximum in our studies), although some animals are found with no obvious proximal cause of death (FDIC, found dead in cage) and some are euthanized due to acutely severe immobility. As with ataxia and clasping, enrofloxacin appears to accelerate disease: enrofloxacin was associated with a significant reduction in survival in *Ndufs4*(-/-)/*MyD88*(Ctl) (***Fig. 5A***). In contrast, *MyD88* loss led to a modest but significant increase in median survival when comparing enrofloxacin treated *Ndufs4*(-/-)*/MyD88*(Ctl) animals (***Fig. 5A***). Maximum survival was also notably increased.

**Figure 5.**
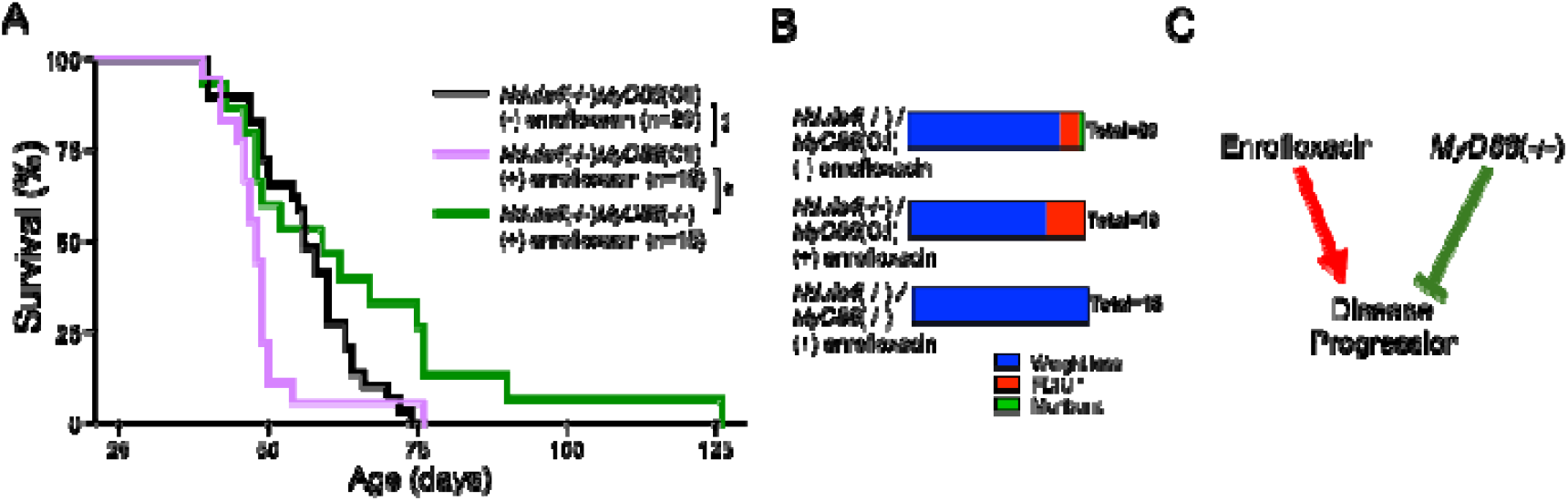
*MyD88* deficiency and enrofloxacin differentially impact survival in *Ndufs4*(-/-) mice. A) Impact of enrofloxacin and *MyD88* disruption on survival in *Nduf4*(-/-) mice (see ***Methods***, introduction). P-values shown indicate pairwise log-rank test comparisons between curves indicated - **p=0.0087, *p<0.013. n’s as indicated. B) Cause of death in mice from A). FDIC – found dead in cage, cause of death unknown. C) Summary of findings. Enrofloxacin treatment modestly accelerates disease progression in the *Ndufs4*(-/-) model, while loss of *MyD88* modestly slows disease progression and extends survival.

Interestingly, *MyD88* deficiency may alter the cause of death in *Ndufs4*(-/-) mice. While a portion of *Ndufs4*(-/-) mice in every other cohort died from unknown causes, all *MyD88* deficient animals were euthanized due to reaching their weight cutoff (***Fig. 5B***). Similarly, enrofloxacin may increase the number of animals that die from unknown causes (found dead in cage, FDIC). Overall, our findings indicate that *MyD88* loss modestly delays disease onset and progression in the *Ndufs4*(-/-), while enrofloxacin modestly accelerates disease (***Fig. 5C***).

## Discussion

### MyD88,TLRs, and Leigh syndrome

Here, we have directly tested, using a genetic approach, whether *MyD88* mediated innate immune signaling downstream of TLRs plays a significant role in the pathogenesis of Leigh syndrome. While we find that while *MyD88* loss appears to modestly delay the onset of neurologic symptoms and extend survival, *MyD88* loss did not alter overall disease pathogenesis in the LS model. *MyD88* deficient *Ndufs4*(-/-) animals still develop the sequelae present in *MyD88* competent *Ndufs4*(-/-) mice, including neurologic disease and the disease-defining neuroinflammatory lesions in the brainstem. This is in contrast to treatment with the CSF1R inhibitor pexidartinib, which ablates multiple immune cell types and appears to fully suppress Leigh syndrome features in this animal model (2). Accordingly, our primary conclusion is that *MyD88*-mediated signaling is not a major driver of Leigh syndrome. Based on current understanding of TLRs, this provides evidence against all among them - except TLR3 - as primary causal drivers of disease.

### Immune modulation likely underlies the benefits of various pre-clinical compounds in Leigh syndrome

*MyD88* deficiency leads to immune suppression beyond disruption of TLR signaling including impaired IL1R and IL-18R signaling, defects in T-cell proliferation, and impaired macrophage responses to IFNγ (26, 33). The modest (albeit statistically significant) delay in disease and extension of survival associated with *MyD88* deficiency is reminiscent of many reports in the *Ndufs4*(-/-) model where treatments somewhat slow or delay symptoms and death but do not appear to greatly alter the overall pathogenesis of Leigh syndrome. Examples include PKC inhibitors, mTOR inhibition (rapamycin), PI3Kγ inhibition, low dose pexidartinib (in contrast to high dose which prevents disease), doxycycline, IFNγ depletion, and others (2, 15, 17, 18, 34-36). Our overall interpretation of these studies and our findings with *MyD88* is that any intervention which has an overall immunosuppressive effect, even modest, is likely to modestly attenuate disease course in the *Ndufs4*(-/-). However, modest immune suppression is not sufficient to impact disease in a robust way as observed with high dose pexidartinib treatment.

This perspective leads to two suggestions for the field going forward: first, statistical significance should not alone be considered sufficient to suggest biological or clinical significance. High dose pexidartinib and hypoxia provide both (2, 37), while *MyD88* deficiency provides only the former. Second, all interventions shown to provide a modest benefit should be considered through the immune perspective. For example, doxycycline was interpreted as acting through suppression of mitochondrial translation but is known to be immune suppressive and was not shown to alter mitochondrial translation *in vivo*, but did have reported immune effects, in the study where it extended *Ndufs4*(-/-) survival (34). Similarly, in our initial work with rapamycin it appeared metabolic effects mediated disease attenuation, but we’ve since shown that the benefits appear to be related to immune suppression (2, 25).

Our understanding of the cell and molecular pathogenesis of Leigh syndrome has progressed substantially over the past decade and a half. We now know that disease is driven by mitochondrial dysfunction in neurons, that disease progression is mediated by the innate immune system, and that CNS disease involves both central macrophages (microglia) and invasion by peripheral macrophages. The identity of the innate immune activating signal is a key unanswered question in the pathobiology of Leigh syndrome. While we interpret our findings as evidence that *MyD88* deficiency is not a primary driver of Leigh syndrome, the importance of this and similar negative results should be emphasized. As our gene expression data demonstrate, individual factors do not need to play a causal role to appear as part of broad immune component upregulation in diseased tissues. Only causal studies, such as the use of knockout animals or robust pharmacologic interventions, can directly test whether candidate pathways are involved in disease. Our findings here rule out *MyD88* as a major causal pathway and allow us to prioritize the next candidate. Given the relatively few known innate immune activation pathways related to mitochondrial components this represents significant progress.

### Synthetic toxicity of enrofloxacin – a notable unexpected finding

Finally, the observation that the enrofloxacin may be toxic in the setting of Leigh syndrome is notable. Enrofloxacin is a fluoroquinolone antibiotic, a class of drugs that inhibit bacterial DNA topoisomerase, gyrase, and helicase, and can impact the homologous mitochondrial enzymes at clinically relevant doses (38-42). Enrofloxacin could impact disease through direct effects on mitochondrial DNA-binding enzymes, via alterations to the microbiome, or by some other process. While Leigh syndrome is known to be highly episodic, the mechanisms driving disease onset and periods of worsening are poorly understood, and relatively little effort has been devoted to exploring drug or environmental interactions in Leigh syndrome (volatile anesthetics are a notable exception (20, 43-45)). These findings emphasize the need for studies exploring drug and environmental interactions with mitochondrial dysfunction.

